# Cross-neutralization of SARS-CoV-2 by HIV-1 specific broadly neutralizing antibodies and polyclonal plasma

**DOI:** 10.1101/2020.12.09.418806

**Authors:** Nitesh Mishra, Sanjeev Kumar, Swarandeep Singh, Tanu Bansal, Nishkarsh Jain, Sumedha Saluja, Jayanth Kumar Palanichamy, Riyaz A. Mir, Subrata Sinha, Kalpana Luthra

## Abstract

Cross-reactive epitopes (CREs) are similar epitopes on viruses that are recognized or neutralized by same antibodies. The S protein of SARS-CoV-2, similar to type I fusion proteins of viruses such as HIV-1 envelope (Env) and influenza hemagglutinin, is heavily glycosylated. Viral Env glycans, though host derived, are distinctly processed and thereby recognized or accommodated during antibody responses. In recent years, highly potent and/or broadly neutralizing human monoclonal antibodies (bnAbs) that are generated in chronic HIV-1 infections have been defined. These bnAbs exhibit atypical features such as extensive somatic hypermutations, long complementary determining region (CDR) lengths, tyrosine sulfation and presence of insertions/deletions, enabling them to effectively neutralize diverse HIV-1 viruses despite extensive variations within the core epitopes they recognize. As some of the HIV-1 bnAbs have evolved to recognize the dense viral glycans and cross-reactive epitopes (CREs), we assessed if these bnAbs cross-react with SARS-CoV-2. Several HIV-1 bnAbs showed cross-reactivity with SARS-CoV-2 while one HIV-1 CD4 binding site bnAb, N6, neutralized SARS-CoV-2. Furthermore, neutralizing plasma antibodies of chronically HIV-1 infected children showed cross neutralizing activity against SARS-CoV-2. Collectively, our observations suggest that human monoclonal antibodies tolerating extensive epitope variability can be leveraged to neutralize pathogens with related antigenic profile.

**Importance:** In the current ongoing COVID-19 pandemic, neutralizing antibodies have been shown to be a critical feature of recovered patients. HIV-1 bnAbs recognize extensively diverse cross-reactive epitopes and tolerate diversity within their core epitope. Given the unique nature of HIV-1 bnAbs and their ability to recognize and/or accommodate viral glycans, we reasoned that the glycan shield of SARS-CoV-2 spike protein can be targeted by HIV-1 specific bnAbs. Herein, we showed that HIV-1 specific antibodies cross-react and neutralize SARS-CoV-2. Understanding cross-reactive neutralization epitopes of antibodies generated in divergent viral infections will provide key evidence for engineering so called super-antibodies (antibodies that can potently neutralize diverse pathogens with similar antigenic features). Such cross-reactive antibodies can provide a blueprint upon which synthetic variants can be generated in the face of future pandemics.

## Introduction

Broadly neutralizing antibodies (bnAbs) targeting the HIV-1 envelope glycoprotein (Env) can neutralize a broad range of circulating HIV-1 isolates and have been called super-antibodies due to their remarkable potency and neutralization breadth (1). As a result of its extensive genetic diversity, HIV-1 is subdivided in multiple clades and circulating recombinant forms (CRFs). A rare subset of HIV-1 infected individuals develops broad and potent antibody responses and have served as potential candidates for the isolation of HIV-1 bnAbs (2, 3). HIV-1 bnAbs take years to develop, have atypical features including long complementarity-determining regions (CDR) loops, high levels of somatic hypermutations (SHMs), presence of insertions and/or deletions (indels), tyrosine sulfation, and develop to tolerate significant alterations in their core epitope (1–3). Notably, V2-apex bnAbs have been shown to exhibit cross-group neutralization activity with viruses derived from HIV-1 group M, N, O and P Envs. Furthermore, they even show cross-neutralization of simian immunodeficiency virus (SIV) isolates (4).

Severe acute respiratory syndrome coronavirus 2 (SARS-CoV-2) emerged in late 2019, rapidly spread across different countries, infecting millions of individuals and has caused a global COVID-19 pandemic (5). The SARS-CoV-2 trimeric spike glycoprotein (S) binds to angiotensin-converting enzyme 2 (ACE2) which leads to host cell entry and fusion (6, 7). Type 1 viral fusion machines, including HIV-1 Env, Influenza hemagglutinin (HA), and SARS-CoV-2 S protein, mediate viral entry driven by structural rearrangements and are trimeric in their pre-fusion and post-fusion state (2, 7, 8). SARS-CoV-2 S protein is covered by host-derived glycans on 66 PNGS on each trimer and site-specific glycan analysis has shown that 28% of glycans on the protein surface are underprocessed oligomannose-type glycans (9). SARS-CoV and HCoV OC43 elicited antibodies have been shown to cross-react with SARS-CoV-2. The Neutralizing antibody (nAb), S309, isolated from memory B-cells of a SARS-CoV infected individual targets a glycan epitope conserved within the Sarbecovirus subgenus (10). Several HIV-1 bnAbs have been shown to penetrate the glycan shield and contact protein residues in Env via their long complementary determining region (CDR) loops and make stabilizing contacts with the surrounding high mannose and complex glycans (11, 12). Several HIV-1 bnAbs recognize glycopeptides and/or cluster of N-linked glycans (1–3). The glycans on HIV-1 Env are highly dynamic and can be occupied by different glycoforms due to glycan processing. The glycan shield covering the HIV-1 Env comprises roughly half its mass and shields approximately 70% of the protein surface with glycosylation occurring on potential N-linked glycosylation sites that vary significantly between infected individuals (18 – 33 PNGS) (13, 14).

Herein, we reasoned that given the unique nature of HIV-1 bnAbs and their ability to recognize and/or accommodate viral glycans, the glycan shield of SARS-CoV-2 spike protein can be targeted by HIV-1 specific bnAbs.

## Results and Discussion

In the past decade, a large panel of bnAbs and non-nAbs targeting diverse epitopes on the HIV-1 Env glycoprotein have been isolated and extensively characterized (reviewed in refs (1–3)). To evaluate the potential cross-reactivity of these antibodies, we first performed binding ELISA of 30 bnAbs and 7 non-nAbs with SARS-CoV-2 S2P_ecto_ protein (pre-fusion stabilized ectodomain construct, 1 – 1208 amino acid residues) and receptor-binding domain (RBD, residues 319 – 541, also called S1^B^ domain). The HIV-1 bnAbs were categorized into five categories based on their epitopes on the viral Env (**figure 1A)**. CR3022, a nAb isolated from a convalescent SARS-CoV patient (15), which has been shown to cross-react with SARS-CoV-2 (7), was used as positive control while two antibodies targeting the envelope glycoprotein of simian immunodeficiency virus were used as negative control. Of the 30 HIV-1 bnAbs tested for binding to both S2P_ecto_ protein and RBD of SARS-CoV-2, 6 bnAbs (VRC07.523LS, NIH45-46 G54W, N6, Z13e1, 2F5 and 4E10) showed significant binding, while one bnAb (CAP256.09) showed moderate binding to only the S2P_ecto_ protein (**figure 1A**). Non-nAbs that target post-fusion and/or open trimeric conformation of HIV-1 Env were unable to bind both SARS-CoV-2 S2P_ecto_ protein and RBD protein (**figure 1B**), suggesting that only pre-fusion state specific antibodies that evolve via extensive somatic hypermutation and affinity maturation in response to repeated exposure to a continuously evolving antigen can cross react with other viruses.

**Figure 1.**
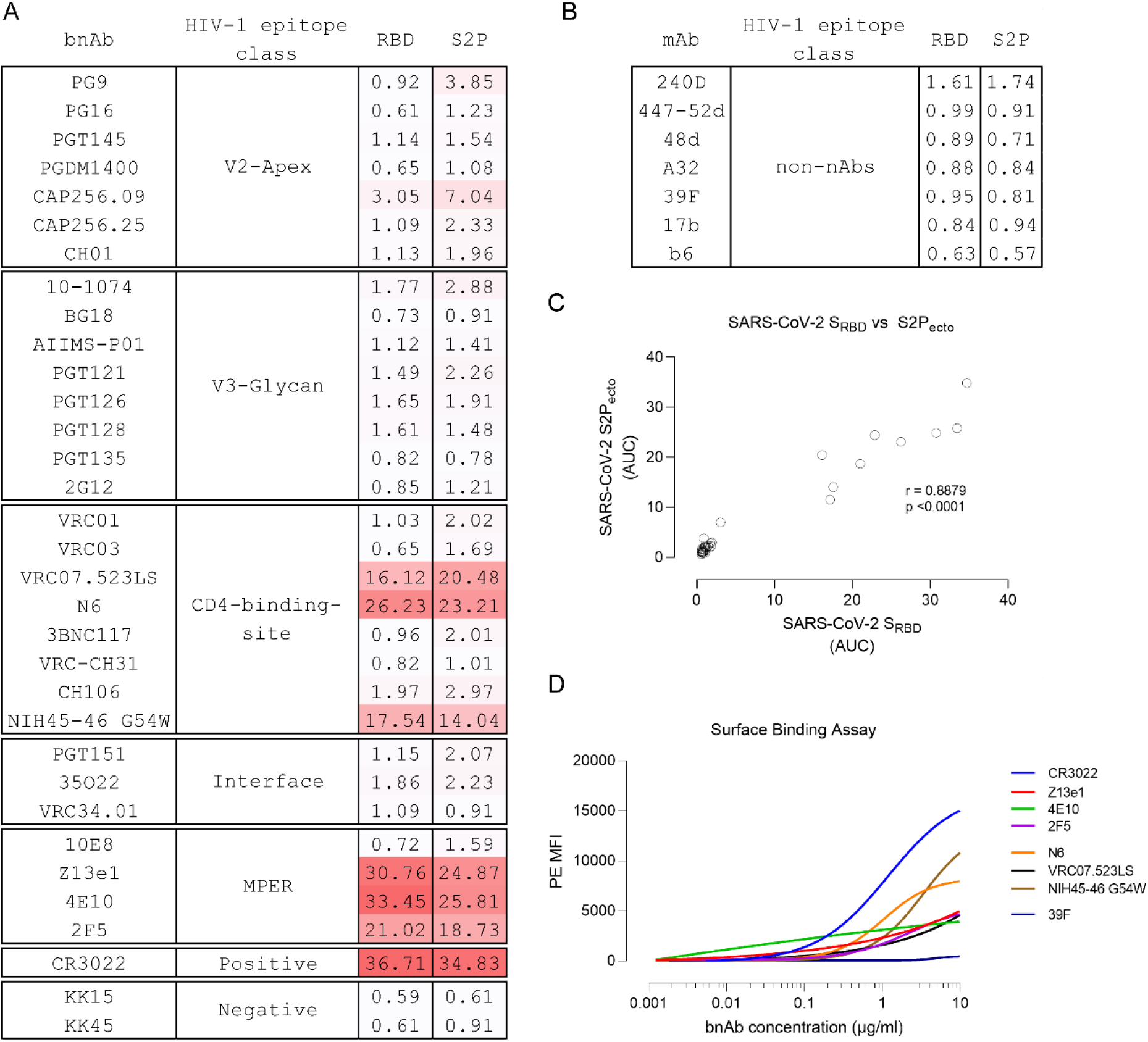
HIV-1 bnAbs cross-react with the receptor binding domain of SARS-CoV-2. (A – B) Cross-reactivity of anti-HIV-1 broadly neutralizing antibodies targeting diverse epitopes on HIV-1 Env and non-neutralizing antibodies were assessed by ELISA using SARS-CoV-2_RBD_ and SARS-CoV-2 S2P_ecto_. CR3022, a SARS-CoV neutralizing antibody, was used as positive control. Two antibodies targeting SIV Env were used as negative control. Area under curve (AUC) of OD_450_ values of a 12-point binding curve (range, 0.0048 to 10 μg/ml) from three independent experiments are shown. (C) Two-tailed Spearman’s correlation was calculated using the area under curve (AUC) values. A significant positive correlation was observed between RBD and S2P_ecto_ (spearman r = 0.8879, p <0.0001). (D) Binding of HIV-1 bnAbs that showed cross-reactivity to S2P and RBD domain of SARS-CoV-2 in ELISA to full-length SARS-CoV-2 S glycoprotein expressed on the surface of HEK293T cells. Average median fluorescence intensity values of a 12-point binding curve (range, 0.0048 to 10 μg/ml) from three independent experiments were used to draw the curve. CR3022, a SARS-CoV neutralizing antibody, was used as positive control.

Though reactivity against RBD was stronger than S2P_ecto_ protein for majority of the bnAbs tested, a significant positive correlation was seen between binding to S2P_ecto_ protein and RBD (**Figure 1C**). All the 6 bnAbs that showed binding reactivity in ELISA exhibited a similar binding profile to the cell surface expressed SARS-CoV-2 S glycoprotein (**figure 1D**). Of the 30 monoclonal antibodies tested herein, bnAbs targeting the membrane proximal external region (MPER) of HIV-1 showed maximum binding to both the S protein and RBD with half-maximal effective concentration (EC_50_) of 0.71 μg/ml and 1.71 μg/ml (Z13e1), 0.048 μg/ml and 2.91 μg/ml (2F5) and 0.79 and 0.33 μg/ml (4E10) μg/ml to the RBD and S2P_ecto_ respectively. VRC07.523LS is an engineered variant of the VRC01 bnAb with higher SHM (16) and while it showed binding to both S2P_ecto_ protein and RBD, both VRC01 and its somatic related clone VRC03 did not show any binding (**figure 2A – B**).

**Figure 2.**
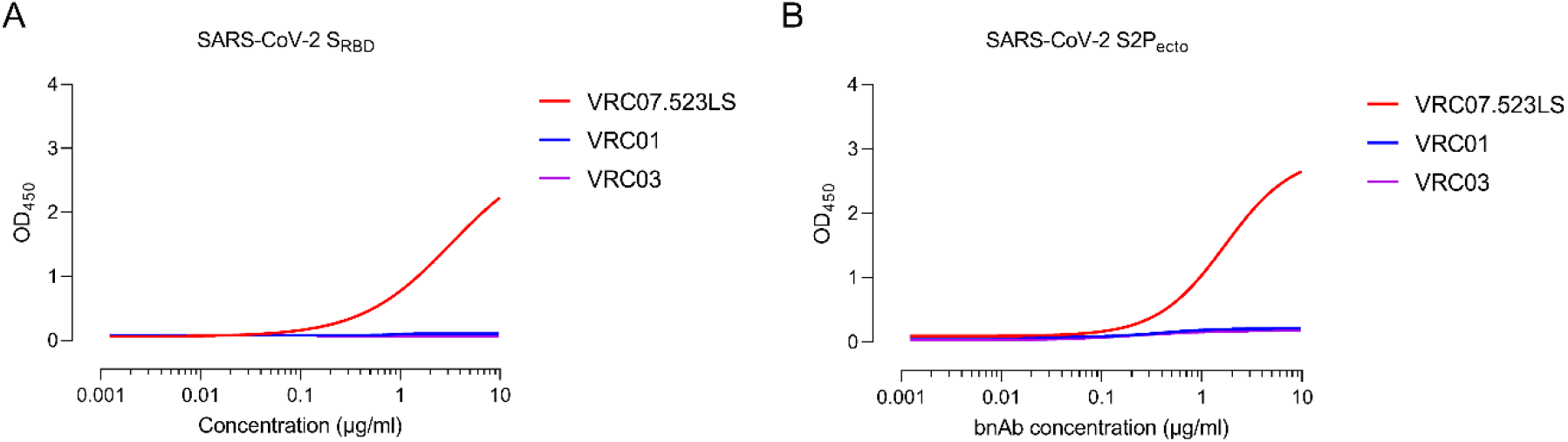
Somatically engineered VRC07.523LS cross-reacts with SARS-CoV-2. Cross-reactivity of anti-HIV-1 broadly neutralizing antibodies, VRC07.523LS, VRC01 and VRC03, targeting the CD4-binding site on HIV-1 Env. Cross-reactivity was assessed by ELISA using (a) SARS-CoV-2_RBD_ and (b) SARS-CoV-2 S2P_ecto_. OD^450^, optical density at 450 nm. OD^450^ values are from a 12-point binding curve (range, 0.0048 to 10 μg/ml).

All 6 HIV-1 bnAbs that showed binding to SARS-CoV-2 S protein and RBD were then tested for their ability to block infection using a HIV-1 pseudovirus based neutralization assay utilizing SARS-CoV-2 spike protein. VSV-G and MLV pseudotyped viruses were used as negative control. Except N6, all remaining five bnAbs failed to neutralize SARS-CoV-2 (**figure 3A**). Though N6 showed neutralization of SARS-CoV-2, it failed to show complete neutralization (maximum percent neutralization of 88% with an IC_50_ of 0.988 μg/ml) and had a moderate affinity of 1.04 × 10^8^ M (**figure 3B**). Furthermore, N6 failed to block RBD binding to soluble ACE2 by ELISA (**figure 3C**), suggesting it recognizes an epitope on RBD outside the ACE2 binding site. It is noteworthy that N6 is a member of the VRC01 class of antibodies that target the CD4bs of HIV-1; it recognizes HIV-1 in an unusual orientation and neutralizes HIV-1 isolates that are typically resistant to other VRC01 class bnAbs (17). Furthermore, it can tolerate absence of key CD4bs antibody contact residues across the length of heavy chain and can tolerate escape mutations that typically provide resistance to HIV-1 from other CD4bs bnAbs. Of note, N6 has an unprecedented degree of somatic hypermutation (31% in heavy and 25% in light chain at the nucleotide level).

**Figure 3.**
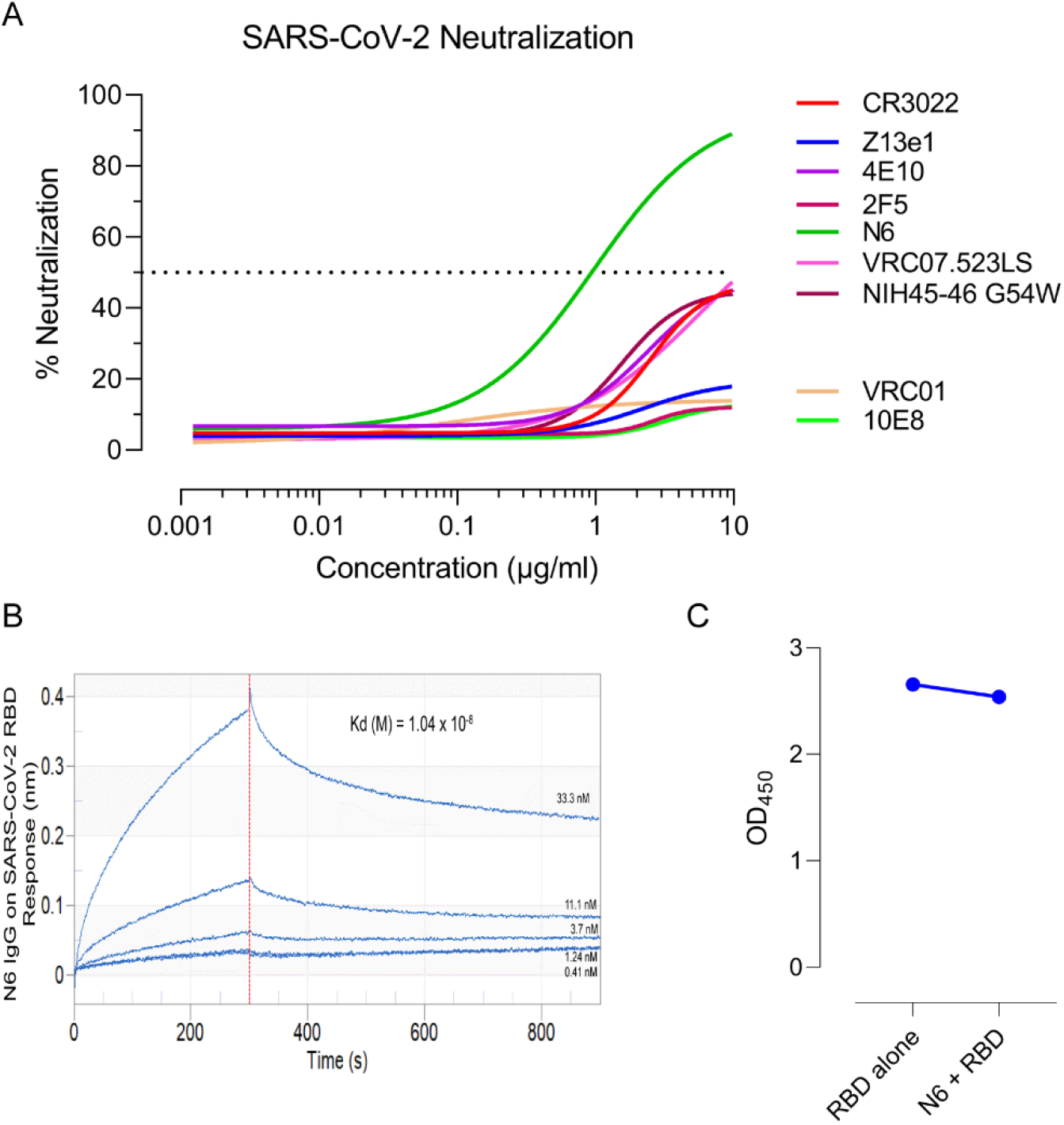
Neutralization of SARS-CoV-2 by HIV-1 bnAbs. (A) **The** bnAbs were tested for neutralization of pseudotyped SARS-CoV-2 virions. Percent neutralization was calculated by assessing relative luminescence units (RLU) in cell lysates of HEK293T-ACE2 cells 48 hours after infection with SARS-CoV-2 pseudoviruses in the presence of increasing amounts of bnAbs (range, 0.0048 to 10 μg/ml). N6, an anti-HIV-1 CD4-binding site bnAb, showed cross-neutralization of SARS-CoV-2. Dotted line corresponds to 50% neutralization. Graphs were plotted using average values (percent neutralization) from three independent experiments. (B) Affinity of N6 against SARS-CoV-2 RBD was measured using biolayer interferometry. (C) Competition ELISA was performed for RBD binding to ACE2 in presence and absence of N6. Average OD^450^ value from three independent experiments are shown.

We next tested plasma antibodies of children with chronic HIV-1 infection for their ability to bind SARS-CoV-2 S2P_ecto_ protein and RBD. Ten children that had shown potent neutralization titre against a 12-virus global panel of HIV-1 isolates from previous studies in our lab were selected (18–20). While all ten children showed significant binding to both S2P_ecto_ and RBD (**figure 4A**), three children showed potent and near-complete neutralization of SARS-CoV-2 pseudoviruses (AIIMS329, AIIMS330, AIIMS346) while two children (AIIMS355 and AIIMS521) showed moderate neutralization of SARS-CoV-2 (**figure 4B – C**). Collectively, our findings highlight the ability of HIV-1 specific bnAbs and polyclonal plasma to cross-react with the newly emerged SARS-CoV-2.

**Figure 4.**
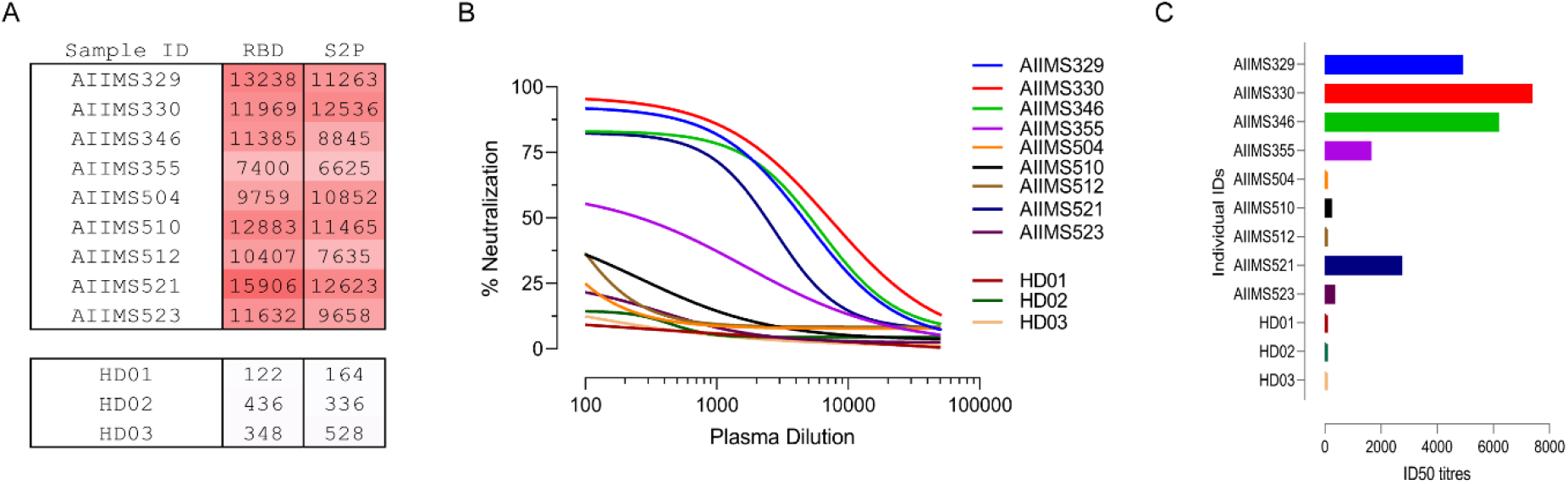
Polyclonal plasma of HIV-1 infected children neutralizes SARS-CoV-2. (A) Cross-reactivity of anti-HIV-1 neutralizing plasma antibodies from ten children with chronic HIV-1 infection against SARS-CoV-2^RBD^ and SARS-CoV-2 S2P^ecto^ was assessed by ELISA. Plasma antibodies from three seronegative healthy donors were used as negative control. Area under the curve (AUC) of OD^450^ values of a 12-point binding curve (range, inverse plasma dilution of 100 to 51200), from three independent experiments are shown. (B) Plasma samples were tested for their neutralization of pseudotyped SARS-CoV-2 virions. Percent neutralization was calculated by assessing relative luminescence units (RLU) in cell lysates of HEK293T-ACE2 cells 48 hours after infection with SARS-CoV-2 pseudoviruses in the presence of increasing dilution of plasma samples (range, inverse plasma dilution of 100 to 51200). (C) Respective ID^50^ (50% inhibitory dilution) for plasma from all ten children are shown.

Neutralizing antibodies engage the host immune system to clear the pathogen or infected cells and are promising candidates for combating emerging viruses (21–23). The RBD of coronaviruses are highly immunogenic and infected individuals typically mount a nAb response (10, 24–28). Given that several HIV-1 bnAbs showed cross-reactivity with RBD of both SARS-CoV and SARS-CoV-2, vaccine efforts should focus inducing antibodies targeting the cross-reactive epitopes on RBD. Most HIV-1 vaccine candidates are in the stage where they typically induce tier 1B or 2 responses against autologous and heterologous viruses in rabbits and non-human primates (29–31). A germline targeting HIV-1 candidate immunogen (eOD-GT8) which was designed to prime VRC01 class CD4bs directed antibodies has been described and the frequencies and affinity of B cells from healthy HIV-1 uninfected individuals recognizing this germline-targeting immunogen showed its suitability as a candidate human vaccine prime (32, 33). Naive B-cells that recognised eOD-GT8 had L-CDR3 sequences that matched several VRC01 class bnAbs, suggesting B-cells with light chain sequences for VRC01 class exist at high frequency. Based on the above observations and the availability of sera from these immunized animals, and findings herein of the ability of HIV-1 CD4bs directed bnAbs to inhibit SARS-CoV and SARS-CoV-2 pseudovirus infection, it is pertinent that the immune sera be tested for binding and neutralization of SARS-CoV-2. Furthermore, detailed structural studies should be taken with N6 to identify its epitope and neutralization determinants, which can be used to engineer its variants as effective SARS-CoV-2 therapeutics.

Understanding cross-reactive neutralization epitopes of antibodies generated in divergent viral infections can provide key evidence for engineering so called super-antibodies (antibodies that can potently neutralize diverse pathogens with similar antigenic features). Such cross-reactive antibodies can provide a blueprint upon which synthetic variants can be generated in the face of future pandemics.

## Methods

### Study design

The current study was designed to assess the cross-reactivity of HIV-1 broadly neutralizing antibodies and plasma antibodies from children with chronic HIV-1 infection against the SARS-CoV-2. The study was approved by the institute ethics committee of All India Institute of Medical Sciences (IEC/NP-295/2011 & IEC/59/08.01.16).

### Cell lines

HEK293T cells for pseudovirus production and generation of 293T-ACE2 cells, and TZM-bl cells for HIV-1 pseudovirus neutralization assay were maintained at 37°C in 5% CO_2_ DMEM containing 10% heat-inactivated FBS (vol/vol), 10mM HEPES, 1mM sodium pyruvate, and 100 U ml^−1^ penicillin/streptomycin. Expi293F cells for recombinant antigen and monoclonal antibody production (Thermo Fisher Scientific, A1452) were maintained at 37°C in 8% CO_2_ in Expi293F expression medium (Thermo Fisher Scientific, A1435102).

### Plasmids

phCMV3 expression plasmids encoding the soluble S2P ectodomain of SARS-CoV (residue 1 – 1190), SARS-CoV-2 (residue 1 – 1208), RBD domain of SARS-CoV (residue 319 – 513) and SARS-CoV-2 RBD (residue 332 – 527) were kindly gifted by Dr. Raiees Andrabi (The Scripps Research Institute). pCR3 expression vectors encoding truncated version of SARS-CoV S protein (residue 1 – 1236) and SARS-CoV-2 S protein (residue 1 – 1254), and pNL4-3ΔEnv-nanoluc were kindly gifted Dr. Paul Bieniasz (The Rockefeller University). CR3022 fab heavy (GenBank: DQ168569.1) and light (GenBank: DQ168570.1) chains were synthesized commercially and subcloned in phCMV3.

### Bacteria

E. coli DH5α, DH10β and STBL3 for propagation of plasmids were cultured at 37°C C (30°C C for STBL3) in LB broth (Sigma-Aldrich) with shaking at 220 rpm.

### Plasma from children with chronic HIV-1 infection

Well-characterized plasma sample from ten children that had shown potent neutralization titre against a 12-virus global panel of HIV-1 isolates from previous studies in our lab were selected.

### Recombinant protein production and purification

SARS-CoV and SARS-CoV-2 ectodomain and RBD constructs were transiently transfected in Expi293F cells at a density of 2 million cells/mL using polyethylenimine and expression plasmids at a molar ratio of 3:1 and purified from clarified transfected culture supernatants 4-days post-transfection using Ni^2+^-NTA affinity chromatography (GE Life Sciences). Proteins were eluted from the column using 250 mmol/L imidazole, dialyzed into phosphate buffered saline (PBS), pH 7.2 and concentrated using Amicon 10-kDa (RBD) and 100-kDa (S2P_ecto_) Amicon ultra-15 centrifugal filter units (EMD Millipore). Protein concentration was determined by the Nanodrop method using the protein molecular weight and molar extinction coefficient as determined by the online ExPASy software (ProtParam).

### Antibody production and purification

The HIV-1 monoclonal antibodies (PGT145, CAP256.25, VRC01, 10-1074, BG18, AIIMS-P01, and PGT151) were expressed by co-transfection of heavy chain and light chain IgG1 plasmids (1:1 molar ratio) in Expi293F cells at a density of 0.8 – 1.2 million cells/mL using PEI-Max (1:3 molar ratio) as the transfection reagent. Five days post-transfection, antibodies were purified from clarified supernatants using protein A beads, eluted with IgG elution buffer and concentrated using 10-kDa Amicon ultra-15 centrifugal filter units (EMD Millipore).

### Binding ELISA

96-well microtiter plates were coated overnight with 2 μg/ml of purified SARS-CoV S2P_ecro_, SARS-CoV-2 S2P_ecto_, SARS-CoV RBD and SARS-CoV-2 RBD. Plates were blocked with 1% BSA for 3 hours. Monoclonal antibodies were added at a starting concentration of 10 μg/ml, with 11-point titration, and incubated for 2 hours at room temperature. Horseradish peroxidase conjugated goat anti-human IgG was used as secondary antibody and TMB substrate was used for color development. Absorbance at 450 nm was measured using a spectrophotometer.

### Generation of 293T-ACE2 cells

VSV-G pseudotyped lentiviruses packaging the human ACE2 were generated by co-transfecting the HEK293T cells with pHAGE6-CMV-ACE2-ZsGreen plasmid and lentiviral helper plasmids (HDM-VSV-G, HDM-Hgpm2, HDM-Tat and CMV-Rev). 48 hours post-transfection, lentiviruses were harvested and used to infect HEK293T cells pre-seeded 24-hours in the presence of 10 μg/ml polybrene. 3-days post-infection, transduced cells were sorted via flow cytometry and maintained as a polyclonal pool of 293T-ACE2 cells in DMEM containing 10% heat-inactivated FBS (vol/vol), 10mM HEPES, 1mM sodium pyruvate, and 100 U ml-1 penicillin/streptomycin at 37°C in 5% CO_2_.

### Viruses

To generate HIV-1 based SARS pseudotyped viral stocks, HEK293T cells were co-transfected with CMV-Luc, RΔ8.2 backbone plasmid, pTMPRSS2 and pSARS-CoV-S_trunc_ or pSARS-CoV-2_trunc_ using polyethylenimine. Six hours post-transfection, cells were washed twice with RPMI and fresh media (10% DMEM) was added. Supernatants containing virions were harvested 48 hours post-transfection, filtered and stored at −80°C C. infectivity of pseudoviruses was determined by titration on 293T-ACE2 cells.

### Neutralization Assays

SARS-CoV and SARS-CoV-2 S protein were co-transfected with an HIV-1 backbone and helper plasmid expressing firefly luciferase and serine protease TMPRSS2 (CMV-Luc, RΔ8.2 backbone plasmid, pTMPRSS2) in 1.25 × 10^5^ HEK293T cells for 48 hours. Post-transfection, culture supernatants were harvested, filtered and stored at −80°C C. For determination of neutralization potential of bnAbs, eight-point titration curves with 2-fold serial dilution starting at 10 μg/ml, were performed. Serially diluted bnAbs were mixed with pseudotyped viruses for 1 hour at 37°C C. pseudovirus/bnAb combinations were then added to 293T-ACE2 cells pre-seeded (24-hours) at 10,000 cells/well. After 48 – 72 hours, supernatant was removed and luminescence was measured on Tecan luminescence plate reader using Bright Glow reagent. The percent infectivity was calculated as ratio of relative luminescence units (RLU) readout in the presence of bnAbs normalized to RLU readout in the absence of mAb. The half maximal inhibitory concentrations (IC50) were determined using 4-parameter logistic regression (GraphPad Prism version 8.3).

### Biolayer interferometry analysis of the SARS-CoV-2 RBD binding affinity with N6 bnAb

Biolayer interferometry was performed using an Octet Red96 instrument (ForteBio, Inc.). A 5 μg/ml concentration of SARS-CoV-2 RBD-His was immobilized on a Ni-NTA coated biosensor surface. The baseline was obtained by measurements taken for 30 s in running buffer (1x PBS, 0.1% BSA and 0.02% Tween-20), and then, the sensors were subjected to association phase immersion for 300 s in wells containing N6 bnAb diluted in running buffer. Then, the sensors were immersed in running buffer for 600 s to measure dissociation. Biosensor was then regenerated by dipping it in EDTA followed by nickel sulfate solution. The mean Kon, Koff and apparent KD values of the SARS-CoV-2 RBD binding affinity for N6 bnAb were calculated from all the binding curves based on their global fit to a 1:1 Langmuir binding model.

### Cell surface binding assay

1.25 × 10^5^ HEK293T cells seeded in a 12-well plate were transiently transfected with 1.25 μg of SARS-CoV-2 S full-length protein using PEI-MAX. 48 hours post-transfection, cells were harvested and per experimental requirement, distributed in 1.5 ml microcentrifuge tubes. For monoclonal antibody staining, 10 μg/ml of antibody was used and titrated 2-fold in staining buffer. 100 μl of primary antibody (HIV-1 specific monoclonals) were added to HEK293T cells expressing SARS-CoV-2 S, and incubated for 30 minutes at room temperature. After washing, 100 μl of 1:500 diluted PE conjugated mouse anti-human IgG Fc was added, and after 30-minute incubation, a total of 50,000 cells were acquired on BD LSRFortessa X20. Data was analyzed using FlowJo software (version v10.6.1).

### Statistics and Reproducibility

All statistical analyses were performed on GraphPad Prism 8.3. A p-value of <0.05 was considered significant. Neutralization assays were performed in triplicates and repeated thrice. Average IC50 values are shown and used for statistical comparisons. Binding ELISAs were performed in duplicates and repeated thrice. Average OD_450_ values were used for plotting curves. Surface binding assay was performed thrice and average PE-MFI (phycoerythrin-median fluorescence intensity) values were used for plotting curves.

## Acknowledgements

We are grateful to Dr. Raiees Andrabi for providing the S2P_ecto_ and RBD constructs for SARS-CoV and SARS-CoV-2, and Dr. Paul Bieniasz for providing the full-length envelope constructs of SARS-CoV, SARS-CoV-2 and pNL4-3ΔEnv-nanoluc. We are thankful to NIH AIDS Reagent program for providing HIV-1 envelope pseudovirus plasmids, bnAbs, non-nAbs and their expression plasmids, and TZM-bl cells, and Neutralizing Antibody Consortium (NAC), IAVI, USA for providing bnAbs. We are thankful to Dr. Michel Nussenzweig for providing 10-1074 and BG18 bnAb expression plasmids.

## Author contributions

N.M conceived and designed the study, performed binding ELISA, neutralization assay, analyzed data, wrote the initial manuscript, revised and finalized the manuscript. Sd.S. and S.K purified and expressed monoclonal antibodies. N.M and S.S expressed and purified recombinant proteins. T.B and N.J performed binding ELISA. S.K edited and revised the manuscript. R.A.M, S.S and K.L conceived and designed the study. K.L conceptualized and designed the study, edited, revised and finalized the manuscript.

## Data Availability

All data required to state the conclusions in the paper are present in the paper and/or the supplementary data. Source data are provided with this paper. Additional information related to the paper, if required, can be requested from the authors.

